# Dexamethasone inhibits the LPS-induced activation of hypoxia-inducible factor 1 alpha (HIF-1α) in macrophages independently of the DUSP1-p38-TTP-HIF-1α regulatory axis

**DOI:** 10.1101/2025.09.19.677372

**Authors:** Chloe Lockwood, Kalbinder K Daley, John D O’Neil, Katherine J Heighes, Sally A Clayton, Andrew R Clark

**Affiliations:** Department of Inflammation and Ageing, School of Infection, Inflammation and Immunology, College of Medicine and Health, University of Birmingham, Edgbaston B15 2TT, United Kingdom; Kennedy Institute of Rheumatology, Old Road Campus, Roosevelt Dr, Headington, Oxford OX3 7FY, United Kingdom; Department of Immunology and Immunotherapy, School of Infection, Inflammation and Immunology, College of Medicine and Health, University of Birmingham, Edgbaston B15 2TT, United Kingdom

## Abstract

The oxygen-sensitive transcription factor HIF-1α (hypoxia-inducible factor 1α) is a master regulator of cellular and organism-level adaptations to decreasing oxygen availability. Under conditions of oxygen sufficiency HIF-1α is constitutively produced and degraded. In response to hypoxia, the degradation pathway is inhibited and HIF-1α protein accumulates via mechanisms that are now well understood. Additional signals combine with the oxygen-sensitive protein degradation pathway to modulate the expression and/or activity of HIF-1α. In the context of the immune system, HIF-1α can be activated under normoxic conditions by a variety of stimuli, including the pro-inflammatory agonist lipopolysaccharide. Here we use a variety of genetic and pharmacological approaches to reveal that lipopolysaccharide-induced HIF-1α accumulation in primary macrophages is dependent on mitogen-activated protein kinase p38. This response is not dependent on modulation of HIF-1α protein stability: instead, it involves the phosphorylation and inactivation of tristetraprolin, an mRNA destabilizing protein that targets *Hif1a* mRNA for degradation. We previously reported that the glucocorticoid dexamethasone inhibits lipopolysaccharide-induced HIF-1α accumulation and metabolic reprograming in primary macrophages. Here we tested and disproved the hypothesis that dexamethasone prevents HIF-1α accumulation by inducing expression of dual specificity phosphatase 1 and thereby inhibiting p38 function. Hence two novel mechanisms critically regulate HIF-1α activation in lipopolysaccharide-treated macrophages: a p38-dependent mechanism that operates at the post-transcriptional level to control *Hif1a* mRNA expression, and a glucocorticoid-sensitive mechanism that operates at the post-translational level to control HIF-1α protein stability. Combined targeting of these two mechanisms may exert therapeutic effects in contexts where HIF-1α contributes to immune-mediated inflammatory pathology.

## INTRODUCTION

The hypoxia-inducible transcription factor HIF-1α plays a central role in cellular adaptation to oxygen insufficiency, coordinating profound metabolic changes to ensure continued generation of ATP when mitochondrial function is impaired (1,2). The mechanisms by which decreasing oxygen availability leads to the activation of HIF-1α are well understood. When oxygen is freely available HIF-1α protein is constitutively produced but then hydroxylated by oxygen-dependent prolyl hydroxylase (PHD) enzymes. Hydroxylated HIF-1α is then recognized and ubiquitinated by the von Hippel Lindau (VHL) ubiquitin ligase complex, and the lysine 48-linked polyubiquitin chains mark HIF-1α for rapid degradation by the proteasome. When cellular oxygen levels decrease, HIF-1α protein accumulates due to impairment of hydroxylation and degradation processes. HIF-1α then dimerizes with its constitutively-expressed binding partner HIF-1β, binds to hypoxia responsive elements (HREs) that usually contain the consensus sequence (A/G)CGTG, and activates transcription. The genes that are regulated in this manner include several mediators of glycolytic metabolism, such as the glucose transporter *Slc2a1* (GLUT1) and the lactate transporter *Slc16a3* (MCT4). The HIF-1α-mediated shift to glycolytic metabolism enables cells to generate ATP independently of oxidative phosphorylation in mitochondria.

HIF-1α can also be activated by the pro-inflammatory bacterial cell wall component lipopolysaccharide (LPS) in myeloid cells, independently of oxygen availability (3-5). LPS-induced HIF-1α activation promotes a shift to glycolytic metabolism regardless of oxygen sufficiency (the Warburg effect), permitting cells to rapidly increase ATP generation. HIF-1α also promotes pro-inflammatory gene expression, migration and antimicrobial functions, which are essential for protection against pathogens (5-9). However, the mechanism(s) by which LPS and other pro-inflammatory factors promote HIF-1α accumulation are not fully understood. Some studies demonstrated that NF-κB (nuclear factor of the kB enhancer) activated transcription of the *Hif1a* gene (10,11). Others have shown that disruption of the mitochondrial tricarboxylic acid cycle leads to accumulation of metabolites that impair PHD function, promoting HIF-1α accumulation at the post-translational level (12,13). Mitogen-activated protein kinase (MAPK) p38 was implicated in the LPS-induced accumulation of HIF-1α protein in mouse neutrophils and human monocyte-derived dendritic cells (14,15). Disruption of the *Dusp1* (dual specificity phosphatase 1) gene enhanced and prolonged the activation of MAPK p38 by LPS in murine macrophages (16-19) and augmented LPS-induced glycolytic metabolism (20,21). One group of investigators found increased expression of HIF-1α protein in LPS-treated *Dusp1*-/- mouse macrophages (19). Another group found no change of HIF-1α expression in LPS-treated *Dusp1*-/- mouse macrophages, and instead implicated MAPK p38-mediated phosphorylation of the glycolysis regulator PFKFB3 (21). MAPK p38 also contributes to the accumulation of HIF-1α in response to hypoxia, ischemia-reperfusion and other oxidative stresses in mouse embryonic fibroblasts (22), keratinocytes (23) and various tumor cell lines (6,24,25). To our knowledge, only one paper has directly addressed the mechanism by which MAPK p38 regulates HIF-1α expression. Kwon and colleagues proposed that MAPK p38-dependent phosphorylation impairs the interaction of hydroxylated HIF-1α with the VHL complex in ischemic pancreatic cancer cells, preventing its degradation by the proteasome (24). This mechanism has not yet been corroborated in other cell types.

We recently reported that the glucocorticoid dexamethasone destabilized HIF-1α protein in primary macrophages, preventing the LPS-induced glycolytic shift (26). Dexamethasone exerts many of its anti-inflammatory effects by increasing the expression of DUSP1 and thereby down-regulating the MAPK p38 pathway (16,27-31). DUSP1 exerts anti-inflammatory effects by modulating the phosphorylation status and activity of the mRNA-destabilizing factor tristetraprolin (TTP) (32,33). MAPK p38-dependent phosphorylation both stabilizes and inactivates TTP. In *Dusp1*-/- mouse macrophages, where LPS-induced p38 activation is dysregulated, TTP accumulates in its phosphorylated and inactive form, accompanied by increased expression of genes that are negatively regulated by TTP, for example *Tnf* and *Ptgs2* (32,34). Several groups have independently reported that TTP destabilizes *Hif1a* mRNA and reduces expression of HIF-1α protein (35-37).

Drawing together these observations, we proposed two distinct hypotheses to explain how dexamethasone inhibits HIF-1α accumulation and glycolytic metabolism in LPS-activated macrophages. Dexamethasone increases the expression of DUSP1 and thereby inhibits MAPK p38 signaling. According to the first hypothesis, this reduction of MAPK p38 activity favours the activation of TTP and destabilization of *Hif1a* mRNA (i.e. post-transcriptional regulation). According to the second hypothesis, lower p38 activity favours VHL-mediated ubiquitination and proteasome-mediated degradation of HIF-1α protein (i.e. post-translational regulation). To test these hypotheses we used various pharmacological and genetic approaches in both primary mouse bone marrow-derived macrophages (BMDM) and primary human monocyte-derived macrophages (MDM).

## RESULTS

MAPK p38 has previously been implicated in the accumulation of HIF-1α protein in response to various stimuli. To test the involvement of MAPK p38 in LPS-induced HIF-1α expression, primary mouse BMDMs were pre-treated with two chemically distinct p38 inhibitors (SB202190 and VX-745) for 30 minutes prior to the addition of LPS. Dexamethasone was used here as a positive control, which we have previously shown to inhibit LPS-induced HIF-1α accumulation (26). The LPS induced accumulation of HIF-1α protein was also significantly impaired by both p38 inhibitors (Fig. 1A, B). To confirm this finding we investigated *Dusp1*-/- BMDMs, in which the absence of an important negative feedback factor results in dysregulation of MAPK p38 signaling (16). Enhanced and prolonged LPS-induced activation of MAPK p38 in *Dusp1*-/- BMDMs was confirmed (Fig. 1C). This dysregulated signaling was accompanied by increased expression of HIF-1α protein, which was particularly evident at the 8 hour time point (Fig. 1C, D). Expression of *Hif1a* mRNA was dependent on MAPK p38 and enhanced in *Dusp1*-/- BMDMs (Fig. 1E). However, *Dusp1*+/+ and *Dusp1*-/- BMDMs did not differ in *Hif1a* transcription (estimated by measurement of primary transcript) (Fig. 1F). The stability of HIF-1α protein also did not differ between *Dusp1*+/+ and *Dusp1*-/- BMDMs, as demonstrated by cycloheximide chases (Fig 1G). These observations suggest that the DUSP1-p38 pathway may regulate HIF-1α accumulation at the level of *Hif1a* mRNA stability rather than transcription or HIF-1α protein stability.

**Figure 1.**
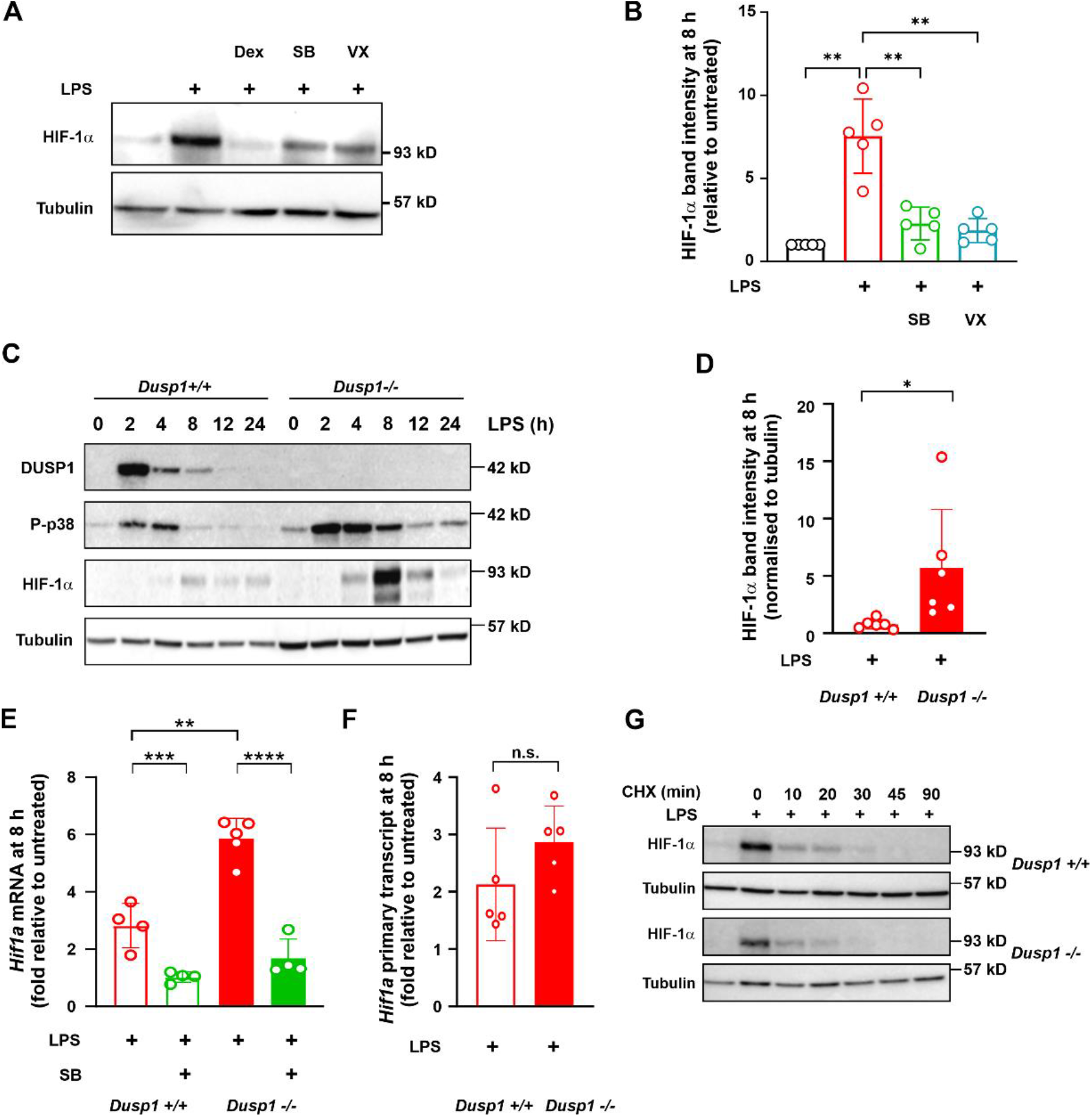
MAPK p38 regulates HIF-α accumulation in LPS-activated mouse BMDMs. Here and throughout, treatment conditions are indicated by different colors and genotypes (where appropriate) are indicated by different shading. **A)** Mouse BMDMs were treated for 8 h with 10ng/mL LPS alone or in combination with 100 nM Dexamethasone, 1 µM SB202190 (SB) or 1 µM VX-745 (VX). HIF-1α and α-tubulin were detected by western blotting. **B)** Quantification of HIF-1α expression (mean ± SEM, normalised to α-tubulin and expressed relative to untreated control; five independent experiments as in A; one way ANOVA with Dunnett’s correction for multiple comparison). **C)** *Dusp1*+/+ and *Dusp1*-/-BMDMs were treated with 10 ng/mL LPS for the indicated times and DUSP1, phospho-p38, HIF-1α and α-tubulin were detected by Western blotting. Representative blots from one of three independent experiments. **D)** Quantification of HIF-1α expression in *Dusp1*+/+ and *Dusp1*-/-BMDMs 8 h after addition of LPS (mean ± SEM, normalised to α-tubulin; six independent experiments; unpaired t test). **E)** *Dusp1*+/+ and *Dusp1*-/-BMDMs were treated with LPS for 8 h in the absence or presence of 1 µM SB202190 (SB). *Hif1a* mRNA was measured by qPCR (normalised to unstimulated control; mean ± SEM of three independent experiments; paired t test within genotype, unpaired t test between genotypes). **F)** *Dusp1*+/+ and *Dusp1*-/-BMDMs were treated with LPS for 8 h and *Hif1a* primary transcript was measured by qPCR (normalised to unstimulated control; mean ± SEM of three independent experiments; unpaired t test). No PCR product was detected if the reverse transcriptase step was omitted, indicating the absence of contaminating genomic DNA. **G)** *Dusp1*+/+ and *Dusp1*-/-BMDMs were treated with LPS for 8 h then cycloheximide (CHX; 5 μg/ml) was added at t = 0 and protein was harvested after the indicated times. HIF-1α and α-tubulin were detected by western blotting. Loading was adjusted for approximately equal HIF-1α band intensity. Representative blot of one of three independent experiments. n.s., p > 0.05; *, p < 0.05; **, p < 0.01.

*Hif1a* mRNA was reported to be targeted for degradation by the RNA-destabilizing protein TTP (35-37). As previously noted (36), the *Hif1a* 3’ UTR sequence is strongly conserved between vertebrate species and contains an adenosine/uridine-rich element with two overlapping consensus TTP binding sites (UUAUUUAUU, highlighted by horizontal lines in Fig. 2A). This arrangement of overlapping motifs is typical of high affinity TTP binding sites and identical to sites that are recognized by TTP in *Tnf* and *Ptgs2* (COX-2) mRNAs (38). MAPK p38 activates its downstream effector kinase MK2, which then phosphorylates TTP at two critical serine residues – Ser52 and Ser178 in mouse. These phosphorylations both stabilize and inactivate TTP protein. Therefore, in *Dusp1*-/- BMDMs, TTP protein accumulates in an inactivate form that is accompanied by increased expression of target mRNAs (32). The accumulation of TTP protein in *Dusp1*-/- BMDMs was confirmed (Fig. 2B). This provides a hypothetical mechanism by which the MAPK p38 pathway could act at the post-transcriptional level to promote *Hif1a* mRNA accumulation; namely p38-dependent phosphorylation and inactivation of TTP, resulting in stabilization of *Hif1a* mRNA. To test this hypothesis we used a knock-in mouse strain in which Ser52 and Ser178 codons of the endogenous mouse TTP locus (formally *Zfp36*) were mutated to alanine codons. In BMDMs derived from this strain (*Zfp36aa/aa*), endogenous TTP protein cannot be phosphorylated and inactivated downstream of the MAPK p38 pathway (39). It therefore behaves as a constitutive negative regulator of target mRNAs, including large numbers of inflammatory mediators (39-42). The LPS-induced expression of *Hif1a* mRNA was impaired in *Zfp36aa/aa* BMDMs, this effect again being most evident at the 8 hour time point (Fig. 2C). Expression of HIF-1α protein was also reduced by approximately 60% in *Zfp36aa/aa* BMDMs (Fig. 2D; also see quantification in Fig. 4A). To test whether the overexpression of *Hif1a* mRNA in *Dusp1*-/- BMDMs was dependent on the phosphorylation of TTP we made use of a double genetically modified strain in which disruption of the *Dusp1* locus was combined with the substitution of Ser52 and Ser178 codons of the *Zfp36* locus. In macrophages derived from this *Dusp1*-/-:*Zfp36aa/aa* strain, the expression of TTP-regulated inflammatory mediators remains low. Although MAPK p38 signaling is elevated, this does not lead to increased gene expression because the critical sites of p38-dependent phosphorylation of TTP are absent (32). *Dusp1* gene disruption caused a statistically significant increase in LPS-induced expression of *Hif1a* mRNA, which could be prevented by combining the knock-out with targeted mutation of TTP Ser52 and Ser178 (Fig. 2E). This demonstrates that DUSP1 regulates the expression of HIF-1α via the phosphorylation of TTP.

**Figure 2.**
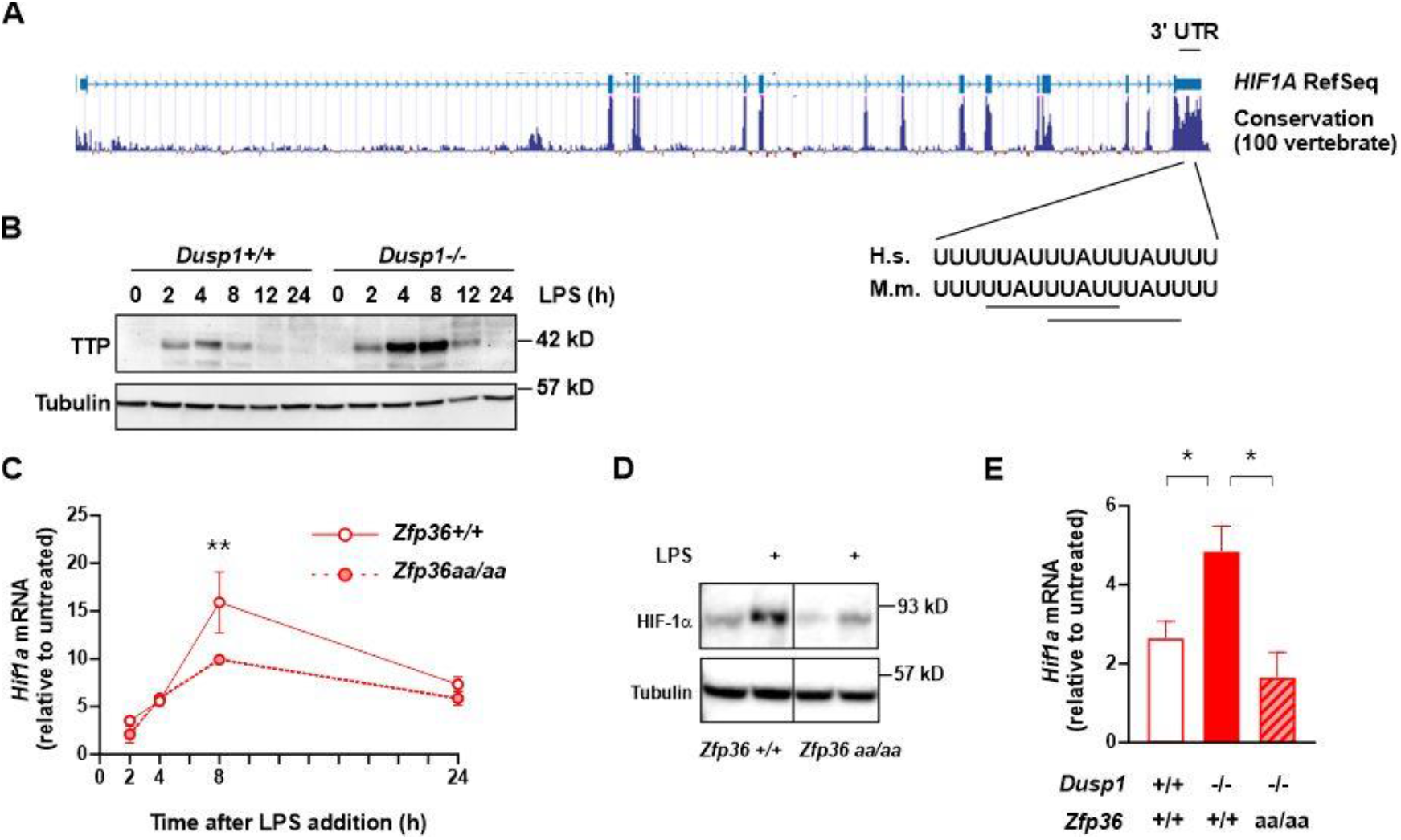
DUSP1 controls HIF-1α expression by modulating the phosphorylation of TTP. **(A)** Evolutionary conservation of the *HIF1A* 3’ UTR between 100 vertebrate species (from UCSC Genome Browser). Conservation of consensus TTP binding sites in mouse *Hif1a* and human *HIF1A* 3’UTRs. **B)** *Dusp1*+/+ and *Dusp1*-/- BMDMs were treated with 10 ng/mL LPS for the indicated times. TTP and α-tubulin were detected by western blotting. Representative of two independent experiments. **C)** *Zfp36*+/+ and *Zfp36aa/aa* BMDMs were treated with 10 ng/mL LPS for the indicated times and *Hif1a* mRNA abundance was measured by qPCR (mean ± SEM fold change relative to untreated control; three independent experiments; two way ANOVA with Dunnett’s correction). **D)** *Zfp36*+/+ and *Zfp36aa/aa* BMDMs were left untreated or stimulated with 10 ng/mL LPS for 8 h. HIF-1α and α-tubulin were detected by western blotting. Representative of three independent experiments. Vertical line represents splicing of different sections of a single gel image. **E)** Wild-type (*Dusp1*+/+:*Zfp36*+/+), DUSP1 knockout (*Dusp1*-/-:*Zfp36*+/+) and double-modified (*Dusp1*-/-:*Zfp36aa/aa*) BMDMs were stimulated with 10 ng/mL LPS for 4 h then *Hif1a* mRNA abundance was measured by qPCR (mean ± SEM fold change relative to untreated control; three independent experiments; one way ANOVA with Dunnett’s correction). *, p < 0.05; **, p < 0.01.

Dexamethasone can exert anti-inflammatory effects by inducing *Dusp1* gene expression and thereby negatively regulating the MAPK p38 pathway (16,27,43). We hypothesised that DUSP1 similarly contributes to the Dex-mediated inhibition of HIF-1α expression and downstream metabolic reprogramming. To test this hypothesis we analysed the effects of Dex on LPS-induced HIF-1α accumulation, glycolysis and expression of HIF-1α-regulated genes of the glycolytic pathway in wild type and *Dusp1*-/- BMDMs. LPS-induced HIF-1α protein accumulation was enhanced in *Dusp1*-/- BMDMs but remained sensitive to Dex (Fig. 3A). LPS-induced lactate secretion and glucose consumption, hallmarks of glycolytic metabolism, were increased in *Dusp1*-/- BMDMs but remained sensitive to Dex (Fig. 3B). LPS-induced expression of the HIF-1α target genes *Slc2a1, Slc16a3* and *Tpi1* was enhanced in *Dusp1*-/- BMDMs but remained sensitive to Dex (Fig. 3C).

**Figure 3.**
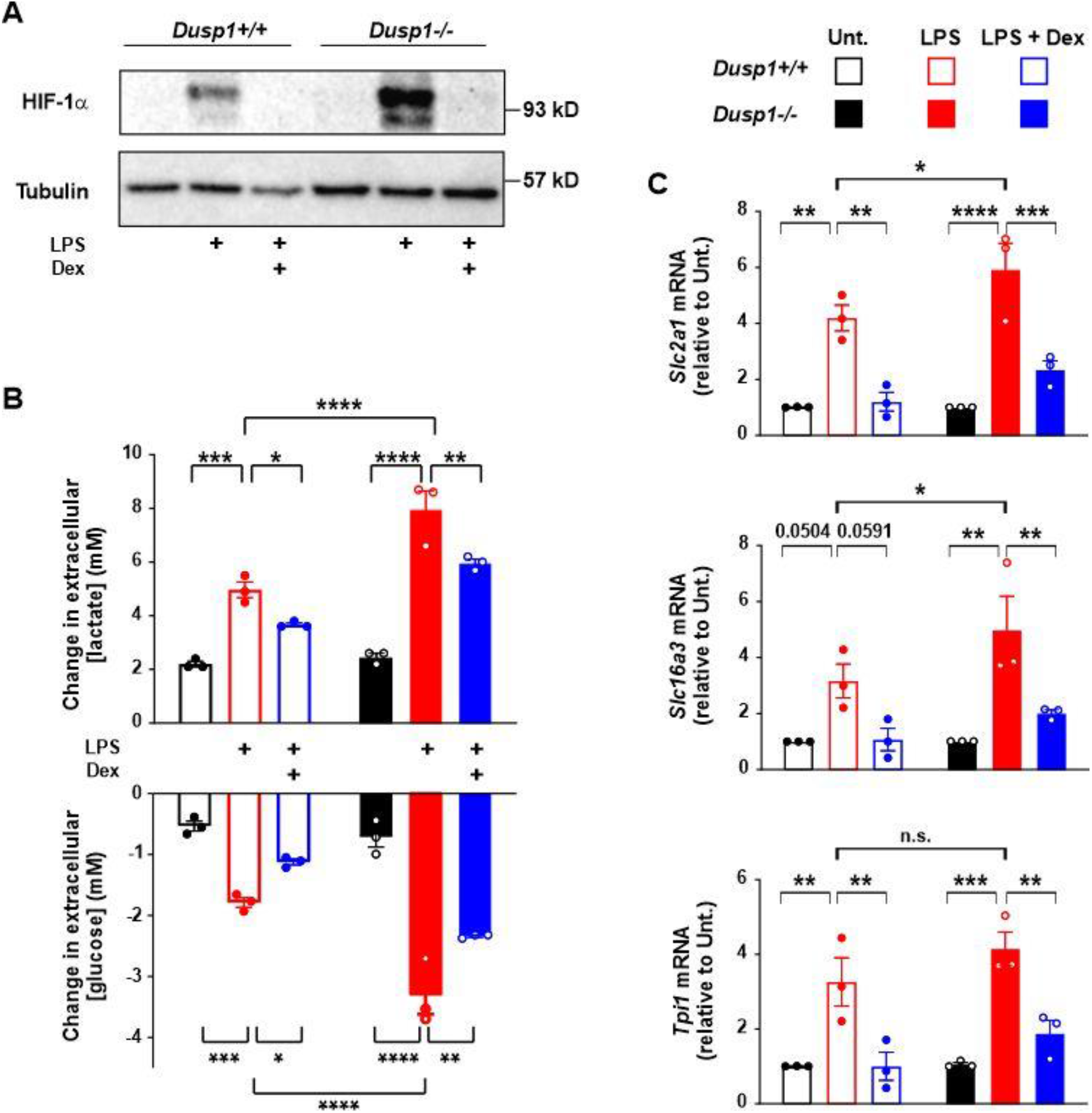
Dexamethasone-mediated inhibition of LPS-induced glycolysis is not dependent on DUSP1. **A)** *Dusp1*+/+ and *Dusp1*-/-BMDMs were treated for 8 h with 10ng/mL LPS alone or in combination with 100nM Dexamethasone. HIF-1α and α-tubulin were detected by western blotting. Representative of six independent experiments. **B)** *Dusp1*+/+ and *Dusp1*-/-BMDMs were treated for 24 h with 10 ng/mL LPS alone or in combination with 100 nM Dexamethasone. Change in extracellular concentration of lactate (upper) or glucose (lower) (mean ± SEM; three independent experiments; two way ANOVA with Sidak’s correction). **C)** Expression of *Slc2a1, Slc16a3* and *Tpi1* mRNA was measured by RT-qPCR after 24 h of treatment as indicated (mean fold change relative to unstimulated control ± SEM; three independent experiments; two way ANOVA with Tukey’s correction). n.s., p > 0.05; *, p < 0.05; **, p < 0.01; ***, p < 0.005; ****, p < 0.001.

Consistent with the lower expression of *Hif1a* mRNA in *Zfp36aa/aa* BMDMs, LPS-induced HIF-1α protein expression was also reduced in *Zfp36aa/aa* BMDMs, but remained sensitive to Dex (Fig. 4A). Interestingly, HIF-1α expression was not significantly affected by the p38 inhibitor SB202190 in *Zfp36aa/aa* BMDMs. Together, these results show that the MAPK p38 pathway regulates HIF-1α expression and upregulation of glycolytic metabolism via the phosphorylation of TTP. Although Dex can induce the expression of DUSP1 and thereby inhibit MAPK p38 in BMDMs (16), the effects of Dex on HIF-1α expression are not dependent on expression of DUSP1 or dephosphorylation of TTP.

If Dex and SB202190 reduce the expression of HIF-1α by distinct mechanisms one might expect the two treatments to exert additive effects. This was difficult to assess in wild-type BMDMs because the powerful inhibitory effect of Dex reduced HIF-1α levels to near undetectable. We therefore used *Dusp1*-/- BMDMs, in which the higher expression of HIF-1α protein allowed us to look for additive effects of the two reagents. The LPS-induced expression of HIF-1α was reduced by either Dex or SB202190 in isolation and eliminated by the two reagents in combination (Fig. 4B, C). Again this supports the argument that Dex impairs HIF-1α expression by a mechanism other than modulation of p38 and TTP.

**Figure 4.**
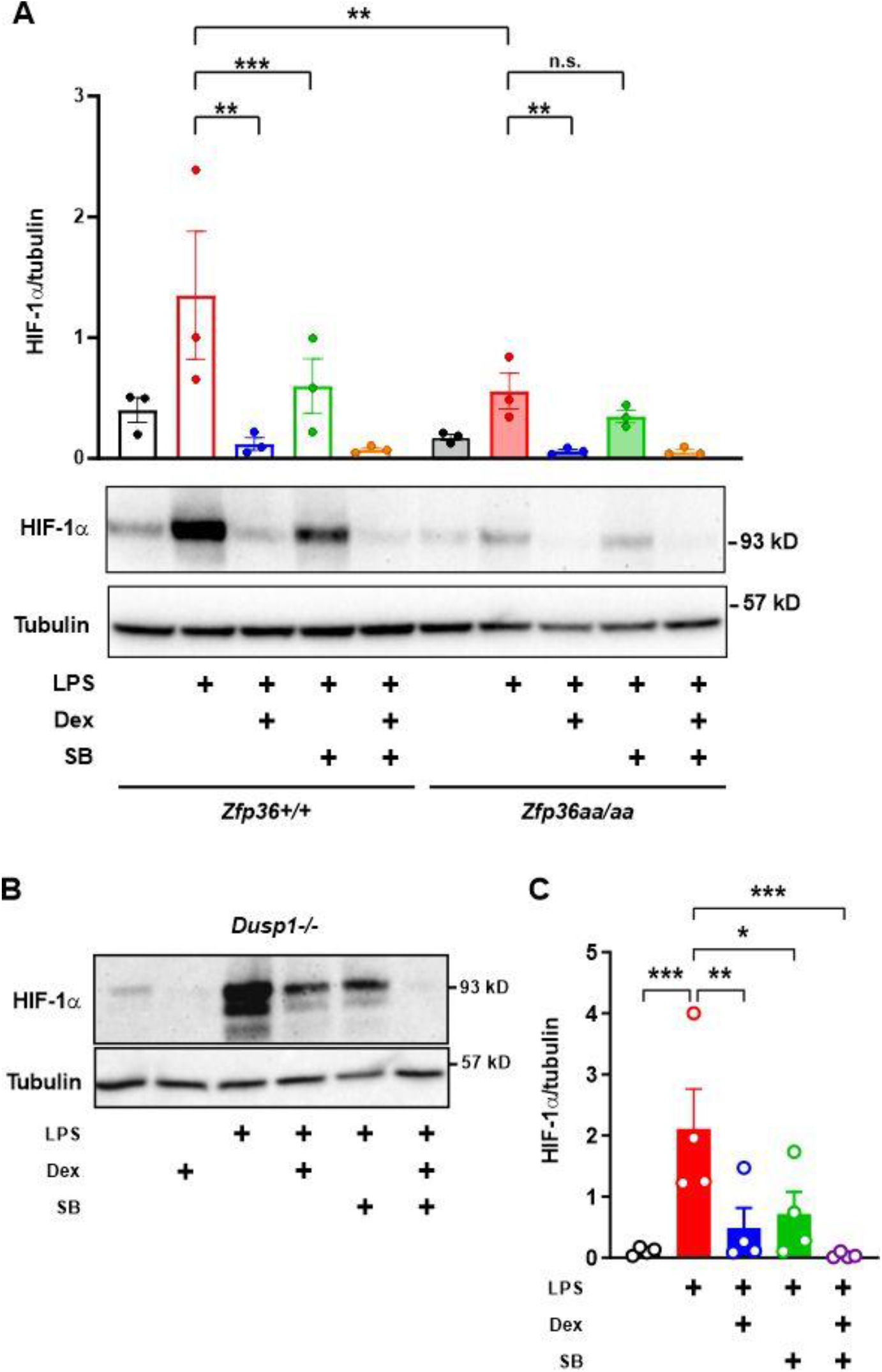
Dexamethasone impairs HIF-1α expression independently of MAPK p38 and TTP phosphorylation. **A)** *Zfp36*+/+ and *Zfp36aa/aa* BMDMs were treated for 8 h with combinations of 10 ng/mL LPS, 100 nM dexamethasone and 1 µM SB202190 as shown. HIF-1α and α-tubulin were detected by western blotting. Graph shows mean ± SEM HIF-1α/α-tubulin ratio from three independent experiments (two way ANOVA with Tukey’s correction). *Dusp1*-/- BMDMs were treated for 8 h with combinations of 10 ng/mL LPS, 100 nM dexamethasone and 1 µM SB202190 as shown. HIF-1α and α-tubulin were detected by western blotting. **C)** Mean ± SEM HIF-1α protein expression (normalised against α-tubulin) in four independent experiments using *Dusp1*-/- BMDMs treated as in B). n.s., p > 0.05; *, p < 0.05; **, p < 0.01; ***, p < 0.005.

We then tested the effects of MAPK p38 inhibitors in human monocyte-derived macrophages (MDMs). SB202190 exerted an inhibitory effect on LPS-induced glycolytic metabolism, which was similar to that caused by Dex (Fig. 5A, B) (26). As in mouse BMDMs, both SB202190 and the structurally unrelated MAPK p38 inhibitor VX-745 reduced the LPS-induced expression of HIF-1α protein in human MDMs (Fig. 5C). As in mouse BMDMs, Dex and SB202190 cooperated with one-another to reduce the expression of HIF-1α protein (Fig. 5D). SB202190 impaired LPS-induced expression of *HIF1A* mRNA, whereas Dex did not do so (Fig. 5E). Both reagents impaired the LPS-induced expression of the HIF-1α target gene *SLC2A1* (Fig. 5F). Dex destabilised HIF-1α protein as previously reported (26), whereas SB202190 did not have this effect under the same conditions (Fig. 5G). Overall, the results presented show that Dex and SB202190 regulate HIF-1α protein accumulation at distinct levels; Dex at the post-translational level and SB202190 at the level of mRNA stability.

**Figure 5.**
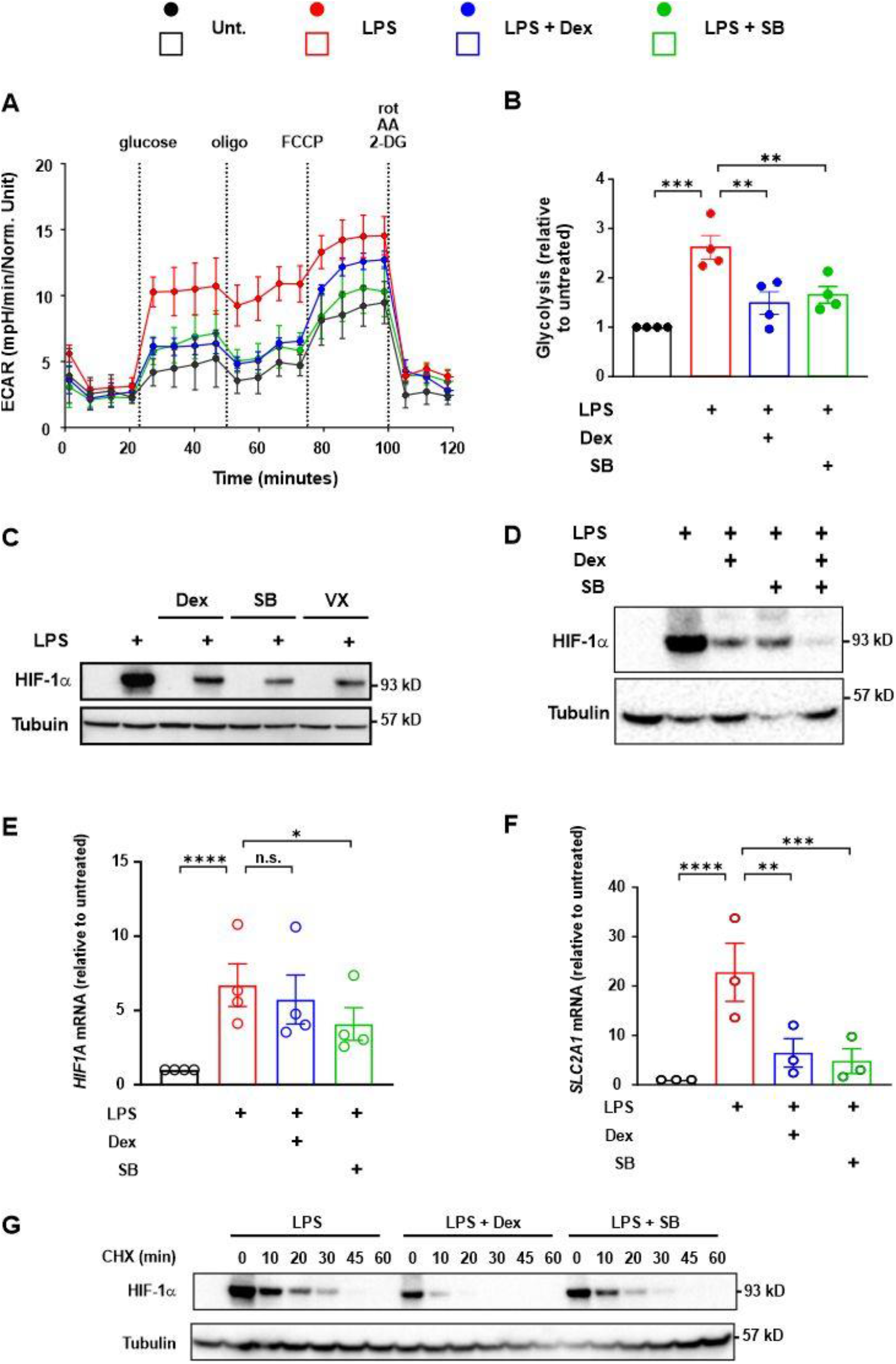
In human MDMs MAPK p38 regulates the LPS-induced glycolytic response at the level of *HIF1A* mRNA expression. **A)** MDMs were treated for 24 h with combinations of 10 ng/mL LPS, 100 nM dexamethasone and 1 µM SB202190 (SB) as shown, then glycolysis was measured by Seahorse XFe96 Mito + Glyco stress test (see materials and methods). **B)** Mean ± SEM fold change in glycolysis relative to unstimulated control from four independent experiments as in A; two way ANOVA with Dunnett’s correction. **C**,**D)** MDMs were treated for 8 h with combinations of 10 ng/mL LPS, 100 nM dexamethasone, 1 µM SB202190 (SB) and 1 µM VX-475 (VX) as shown. HIF-1α and α-tubulin were detected by western blotting. Representative of three (C) or four (D) independent experiments. **E**,**F)** MDMs were treated with 10 ng/mL LPS ± 100 nM dexamethasone or 1 µM SB202190 (SB) for 8 h (peak of *HIF1A* expression) or 12 h (peak of *SLC2A1* expression). Fold change ± SEM of *HIF1A* (E) or *SLC2A1* (F) mRNA abundance relative to unstimulated control; three independent experiments; one way ANOVA with Dunnett’s correction. **G)** MDMs were stimulated for 8 h with 10 ng/mL LPS in the absence or presence of 100 nM dexamethasone or 1 µM SB202190 (SB). Cycloheximide (CHX) was added at t = 0 and cells were harvested at the time points shown. HIF-1α and α-tubulin were detected by western blotting. Representative of two independent experiments. n.s., p > 0.05; *, p < 0.05; **, p < 0.01; ***, p < 0.005; ****, p < 0.001.

## DISCUSSION

The MAPK p38 signaling pathway positively regulates glycolysis in myeloid cells (14,15,20,21). As a master regulator of glycolysis, HIF-1α is a prime candidate mediator of this effect. Indeed, several papers have described regulation of HIF-1α function or expression by the MAPK p38 signaling pathway in different contexts (6,14,15,19,22-25,36,44-46) but few have addressed sites or mechanisms of action. A selective MAPK p38 inhibitor impaired the hypoxia-induced activation of an HRE reporter or a reporter controlled by a Gal4-HIF-1α transcriptional activation domain (TAD) fusion protein in a liver cancer cell line (46). The same inhibitor did not affect HIF-1α protein expression, suggesting that MAPK p38 directly or indirectly regulates function of the HIF-1α TAD. We cannot formally rule out that MAPK p38 also regulates HIF-1α TAD function in macrophages. However, MAPK p38-mediated regulation of HIF-1α expression levels in these cells may be sufficient to explain downstream changes in gene expression and metabolism. Others reported that MAPK p38-mediated phosphorylation of HIF-1α impaired its interaction with VHL and thereby promoted HIF-1α protein stabilization in a pancreatic tumor cell line (24). This mechanism does not appear to operate in macrophages. Hyperactivation of MAPK p38 in *Dusp1*-/- mouse BMDMs was accompanied by increased expression of HIF-1α protein (Fig. 1C, 1D) but no increase in the stability of HIF-1α protein (Fig. 1G). Likewise, HIF-1α protein stability was not affected by a MAPK p38 inhibitor in human MDMs (Fig. 5G).

Based on experiments in THP-1 myeloid leukemia cells, Fähling and colleagues concluded that MAPK p38 controls HIF-1α expression via the phosphorylation of TTP (36). It was important to confirm these findings because THP-1 cells are oncogenically transformed and differ from primary macrophages in several respects (47,48). Using a combination of pharmacological and genetic approaches in primary human and mouse macrophages, here we conclusively demonstrate that MAPK p38 controls HIF-1α expression by modulating the phosphorylation and activity of TTP. Elevated MAPK p38 signaling in *Dusp1*-/- BMDMs was accompanied by increased expression of both *Hif1a* mRNA and HIF-1α protein (Fig. 1C, 1D, 1E). This increased expression could be reversed by MAPK p38 inhibition (Fig. 4B, 4C) or by combining *Dusp1* gene disruption with targeted mutagenesis of the *Zfp36* gene, preventing MAPK p38-mediated inactivation of TTP (Fig. 2E). In TTP mutant (*Zfp36aa/aa*) BMDMs the LPS-induced expression of both *Hif1a* mRNA and HIF-1α protein was impaired (Fig. 2C, Fig. 4A). Furthermore HIF-1α protein expression was no longer sensitive to MAPK p38 inhibition in these cells (Fig. 4A), indicating that MAPK p38 regulates HIF-1α expression principally via the phosphorylation of TTP. Collectively, these findings establish the existence of a DUSP1-p38-TTP-HIF-1α regulatory axis (Fig. 6), confirming and extending previous observations (36). This post-transcriptional mechanism of HIF-1α regulation may be relevant in contexts other than macrophages. For example, tumor-suppressive functions of TTP (49) may involve impairment of HIF-1α accumulation, in addition to direct targeting of mRNAs that encode mediators of glycolysis (50,51). We speculate that, conversely, pro-oncogenic functions of the MAPK p38 pathway (52,53) involve the inactivation of TTP, promotion of HIF-1α accumulation and consequent metabolic adaptation to tumor microenvironments.

**Figure 6.**
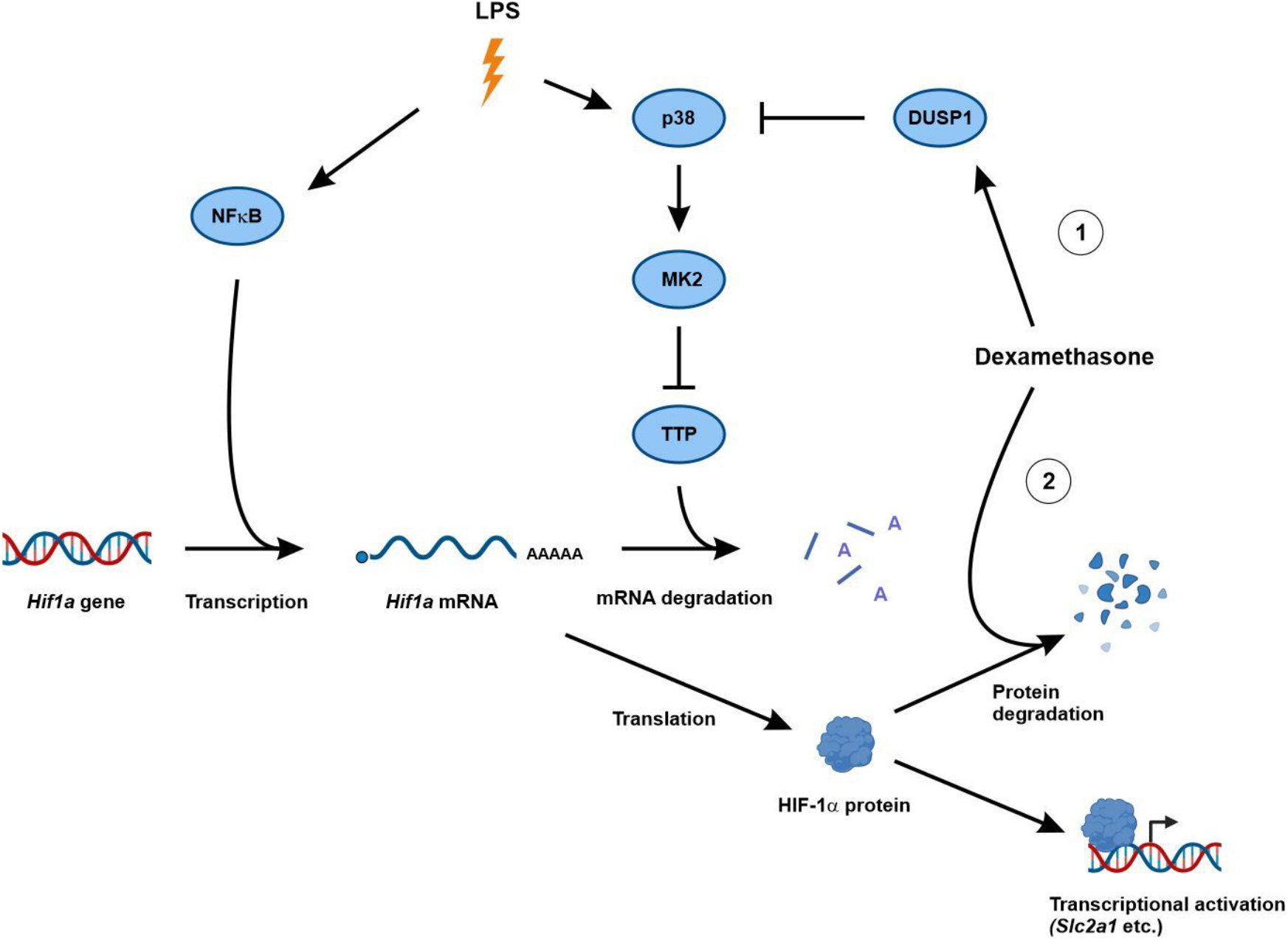
Regulation of LPS-induced HIF-1α expression by dexamethasone. LPS promotes accumulation of *Hif1a* mRNA by activating transcription (e.g. via NF-κB) and by inactivating the mRNA destabilizing factor TTP (via p38-mediated activation of MK2 and MK2-mediated phosphorylation of TTP Ser52 and Ser178). DUSP1 negatively regulates *Hif1a* mRNA accumulation by preventing the phosphorylation and inactivation of TTP. 1) Although dexamethasone can induce DUSP1 expression, this mechanism is dispensible for inhibition of LPS-induced HIF-1α accumulation. 2) Dexamethasone promotes destabilization of HIF-1α protein via a mechanism that remains to be identified.

The glucocorticoid dexamethasone inhibits HIF-1α accumulation and HIF-1α-directed glycolysis in LPS-treated macrophages (26). We hypothesized that dexamethasone exerts this metabolic effect by activating the well-known glucocorticoid target gene *Dusp1* and thereby inhibiting MAPK p38 function (16,30,54,55). Several pieces of evidence contradicted this hypothesis. In MDMs, a MAPK p38 inhibitor significantly reduced the expression of *Hif1a* mRNA in response to LPS treatment, whereas dexamethasone did not. Dexamethasone destabilized HIF-1α protein in LPS-treated macrophages, whereas a MAPK p38 inhibitor did not. Dexamethasone and the kinase inhibitor cooperated to reduce HIF-1α protein expression in both mouse and human macrophages, supporting the argument that they operate at different levels. Most compellingly, in *Dusp1*-/- BMDMs dexamethasone was still able to inhibit the LPS-induced accumulation of HIF-1α protein, expression of HIF-1α-dependent metabolic genes and glycolytic reprograming (Fig. 3). The mechanism(s) by which dexamethasone inhibits LPS-induced HIF-1α expression and glycolytic switch remain(s) to be identified.

The physiological functions of HIF-1α are varied and context-dependent, complicating the question of therapeutic targeting of this transcription factor. HIF-1α contributes to the metabolic reprograming that supports tumor survival and growth in hypoxic environments, therefore inhibitors are actively sought as anti-cancer drugs (56). However, the first HIF-targeting drugs to be licensed were prolyl hydroxylase inhibitors that promote accumulation of HIFs, and are used in the treatment of anemia in chronic kidney disease. HIF-1α has protective, homeostatic functions in the gastrointestinal tract, where its activation by commensal microbes contributes to the maintenance of gut barrier integrity (57). On the other hand, HIF-1α is thought to contribute to pathogenesis of chronic immune-mediated inflammatory diseases such as rheumatoid arthritis (58) and cardiovascular disease (59). In the context of host defence the effects of HIF-1α are two-edged. Several reports have described pathogenic roles in severe COVID-19 (60-62). Yet HIF-1α is essential for defences against intracellular bacteria (6,63-65). Even here, this protective role may be dependent on timing: HIF-1α is protective against early infection of macrophages by *Mycobacterium tuberculosis*, but drives metabolic changes that support prolonged infection (66).

Where there is strong evidence for a pathogenic role of HIF-1α, our findings suggest that combinations of glucocorticoids with MAPK p38 inhibitors may be effective. Whilst glucocorticoids have powerful anti-inflammatory effects, their prolonged use is associated with many side effects having variable severity (67). In principle, combination with other drugs may allow glucocorticoid sparing and mitigation of such side effects. MAPK p38 clearly contributes to the pathogenesis of many immune-mediated inflammatory diseases, yet the deployment of selective inhibitors as anti-inflammatory drugs has proven difficult. In several clinical trials of MAPK p38 inhibitors, therapeutic anti-inflammatory effects were transient and accompanied by side effects that were likely mechanism-related (68). Combination with glucocorticoids could permit therapeutic effects to be achieved with lower doses of inhibitors, minimizing both side effects and tachyphylaxis. These concepts remain to be explored.

## EXPERIMENTAL PROCEDURES

### Macrophage isolation and culture

The generation of *Zfp36aa/aa* mouse strain was described previously (39). Acquisition of *Dusp1*-/- line and the double-targeted *Dusp1-/-Zfp36aa/aa* line was also described previously (32). Mice were housed at the University of Birmingham Biomedical Services unit. All maintenance and procedures were carried out according to the Home Office guidelines and approved by the University of Birmingham Animal Welfare and Ethical Review Board.

Mice were sacrificed by cervical dislocation and hind legs were removed. For bone marrow isolation: femur and tibiae were cleaned of muscle tissue, cut at each end and centrifuged. Extracted bone marrow was placed in RPMI 1640 + L-glutamine (Gibco #21875034) supplemented with 10% heat-inactivated FCS (Sigma Aldrich #F2442) and 50ng/mL M-CSF (PeproTech #300-25) for 7 days to differentiate bone-marrow derived macrophages (BMDMs).

Blood from anonymous healthy human donors was obtained as leukapheresis cones from the NHS Blood and Transplant Service (ethical approval ERN_16-0191). Monocytes were isolated using Ficoll density gradient centrifugation and negative selection with the RosetteSep Human Monocyte Enrichment Cocktail (STEMCELL #15068). Isolated monocytes were placed in RPMI 1640 + L-glutamine (Gibco #21875034) supplemented with 5% heat-inactivated FCS (LabTech #80837) and 50ng/mL M-CSF (PeproTech #300-25) for 7 days to differentiate monocyte derived macrophages (MDMs). We have previously shown that the above methods of macrophage generation yielded populations of primary macrophages that were >95% pure (69).

For stimulations, macrophages were seeded at 1 × 10^6^ cells/mL (mouse) or 0.5 × 10^6^ cells/mL (human) in 12-well or 6-well tissue culture-treated plates. Stimulation medium was prepared with final concentrations of 10ng/mL LPS (E.coli, Serotype EK100 (Ra) (TLRgrade^TM^) – Enzo Life Sciences #ALX-581-010-L002) with/without 100nM dexamethasone (Sigma Aldrich #D18893), or 1µM VX-745 (Stratech A8686-APE) and/or 1µM SB202190 (Stratech A1632-APE). Dexamethasone, VX-745 and SB202190 were dissolved in DMSO – DMSO concentrations were matched across all conditions. For MAPK p38 inhibition experiments cells were pre-treated with VX-745 or SB202190 or vehicle for 30 minutes prior to LPS addition to ensure effective kinase inhibition.

### Lactate/glucose measurements

Lactate and glucose concentrations from BMDM cell cultures were measured using the Nova Stat Profile Prime cell culture analyser. Glucose consumption was calculated by subtracting the glucose concentration of RPMI 1640 (11.11mM) from values obtained from conditioned medium samples.

### Seahorse metabolic flux assays

MDMs were seeded at 50,000 cells/well in Agilent Seahorse XFe96 cell culture microplates and left to adhere overnight. As described (70), we used a combined version of Mito and Glyco stress tests. Seahorse XF RPMI medium, pH 7.4 (Agilent #103576-100) was supplemented with 2 mM L-glutamine (Sigma Aldrich #G7513). The following injection protocol was used (final assay concentrations): (A) D-glucose (10mM, Sigma Aldrich #G7021), (B) oligomycin A (1μM, Cayman Chemical Company #11342), (C) carbonyl cyanide p-trifluoromethoxyphenylhydrazone (FCCP) (5μM, Cayman Chemical Company #15218) + sodium pyruvate (1 mM, Sigma Aldrich #P5280), and (D) rotenone (100nM, Cayman Chemical Company #83-79-4) + antimycin A (1μM, Sigma Aldrich #A8674) + 2-deoxy d-glucose (20mM, Sigma Aldrich #D8375).

For Seahorse normalisation, immediately after assay completion cells were incubated with 1µM calcein-AM viability dye (eBioscience, 65-0853-78) in PBS for 30 min at 37°C. Fluorescence was measured using a plate reader (excitation, 490nm; emission, 515 nm) and the viable cell count ratio was calculated.

### Western Blotting

Cells were lysed directly into 2X XT Sample buffer (Bio-Rad #1610791) and 1X XT Reducing Agent (Bio-Rad #1610792) and passed through a QIAshredder column to remove genomic DNA (QIAGEN, #79656). Samples were then heated to 95°C for 5 min and an equal volume of each sample was loaded onto an XT Bis-Tris protein gel (Bio-Rad). XT MES running buffer (Bio-Rad, #1610789) was used to run the gels for 90 min at 100V. Protein was transferred onto Bio-Rad Trans-Blot polyvinylidene difluoride (PVDF) membranes (Bio-Rad, #1704157) using the Bio-Rad Trans-Blot Turbo transfer system. The desired primary antibodies from **Table 1** were incubated overnight at 4°C. HRP-conjugated secondary antibodies were applied for 1h at room temperature (Cell Signaling Technologies #7074, #7076). Blots were imaged using Clarity Enhanced Chemiluminescence substrate (Bio-Rad #1705061) and the ChemiDoc MP Imaging System (Bio-Rad). Blots were quantified via densitometry using Image J Fiji.

**Table 1.**
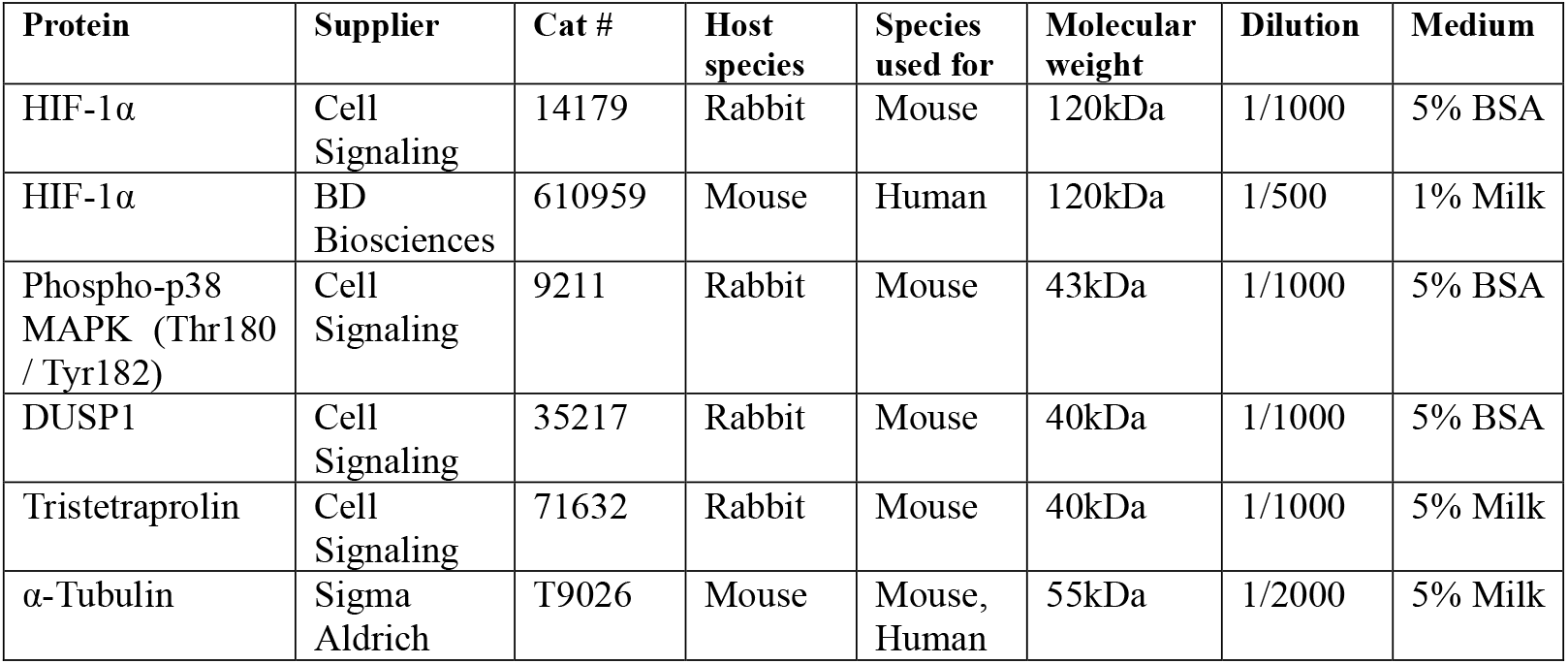
Antibodies used for Western Blotting

### RT-qPCR

RNA was isolated using Norgen Total RNA Purification Plus kits (Geneflow, P4-0016) according to the manufacturer’s instructions. RNA concentrations were quantified using Thermofisher Nanodrop and only samples with A260:A280 (>1.8) and A260:A230 (>1.0) ratios were used for cDNA synthesis. iScript Reverse Transcriptase (Bio-Rad #1708891) was used to synthesize cDNA from 250ng or 500ng RNA/reaction. Gene expression was measured by reverse transcription-quantitative polymerase chain reaction (RT-qPCR) using the Bio-Rad CFX384 system.

SYBR TB Green Premix Ex Taq (Takara, #RR820W) and primers supplied by Sigma Aldrich (**Table 2**) were used. Either *UBC* (human) or *Rpl13a* (mouse) were used to normalise mRNA measurements by the 2^−ΔΔCt^ method.

**Table 2.**
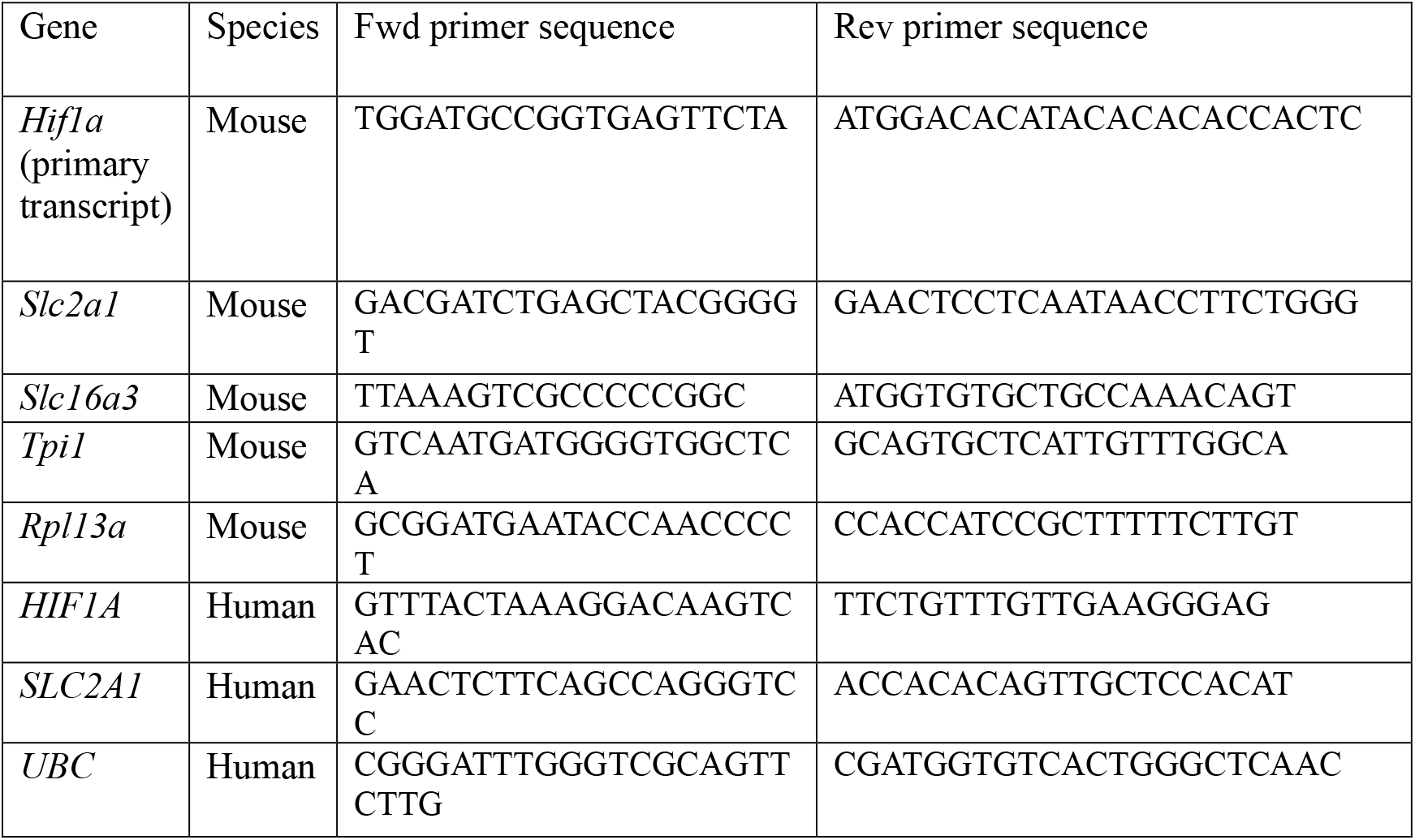
RT-qPCR primer sequences for human and mouse

To measure mature mouse *Hif1a* gene expression a TaqMan gene expression assay (Thermofisher, assay ID: Mm00468869_m1) was used and normalised to Rpl13a (Thermofisher, assay ID: Mm05910660_g1). The TaqMan assay utilised Applied Biosystem’s TaqMan Gene Expression Master Mix (#4369016).

## DATA AVAILABILITY

All relevant data are included within the manuscript.

## ACKNOWLEDGEMENTS

This work was funded by Versus Arthritis Programme Grant 21802 to A.R.C. C.L was funded by a PhD studentship as part of Research into Inflammatory Arthritis Centre Versus Arthritis (grant 22072). S.C. is supported by a Wellcome Trust Early-Career Award (300104/Z/23/Z).

